# Genetic basis underlying connection between hyperglycemia and dyslipidemia in *Apoe*-deficient mice

**DOI:** 10.1101/021287

**Authors:** Qian Wang, Andrew T. Grainger, Ani Manichaikul, Emily Farber, Suna Onengut-Gumuscu, Weibin Shi

**Affiliations:** Departments of Radiology and Medical Imaging (QW, WS) and of Biochemistry & Molecular Genetics (ATG, WS), Center for Public Health and Genomics (AM, EF, SOG), University of Virginia, Charlottesville, VA 22908

## Abstract

Individuals with dyslipidemia often develop type 2 diabetes, and diabetic patients often have dyslipidemia. It remains to be determined whether there are genetic connections between the 2 disorders. A female F_2_ cohort, generated from BALB/cJ (BALB) and SM/J (SM) *Apoe*-deficient (*Apoe*^−/−^) strains, was fed a Western diet for 12 weeks. Fasting plasma glucose and lipid levels were measured before and after Western diet feeding. 144 genetic markers across the entire genome were used for analysis. One significant QTL on chromosome 9, named *Bglu17* [26.4 cM, logarithm of odds ratio (LOD): 5.4], and 3 suggestive QTLs were identified for fasting glucose levels. The suggestive QTL near the proximal end of chromosome 9 (2.4 cM, LOD: 3.12) was detected when mice were fed chow or Western diet and named *Bglu16*. *Bglu17* coincided with a significant QTL for HDL and a suggestive QTL for non-HDL cholesterol levels. Plasma glucose levels were inversely correlated with HDL but positively correlated with non-HDL cholesterol levels in F_2_ mice fed either diet. A significant correlation between fasting glucose and triglyceride levels was observed on the Western but not chow diet. Haplotype analysis revealed that “lipid genes” *Sik3* and *Apoc3* were probable candidates for *Bglu17.* We have identified multiple QTLs for fasting glucose and lipid levels. The colocalization of QTLs for both phenotypes and the sharing of potential causal genes suggest that dyslipidemia and type 2 diabetes are genetically connected.

**Article Summary:** Patients with dyslipidemia often develop type 2 diabetes, and diabetic patients often have dyslipidemia. It remains unknown whether there are genetic connections between the 2 disorders. Using a female F_2_ cohort derived from BALB/cJ and SM/J *Apoe*-deficient mice, we identified one significant QTL on chromosome 9, named *Bglu17*, and 3 suggestive QTLs were identified for fasting glucose levels. *Bglu17* coincided with a significant QTL for HDL and a suggestive QTL for non-HDL levels. Plasma glucose levels were significantly correlated with HDL and non-HDL levels in F_2_ mice. Haplotype analysis revealed *Sik3* and *Apoc3* were probable candidates for both QTLs.

## Introduction

Individuals with dyslipidemia have an increased risk of developing type 2 diabetes (T2D), and diabetic patients often have dyslipidemia, which includes elevations in plasma triglyceride and LDL cholesterol levels and reductions in HDL cholesterol levels (Li *et al.* 2014a). Part of the increased diabetic risk associated with dyslipidemia is probably due to genetic variations that influence both lipoprotein homeostasis and the development of T2D. Indeed, a few rare gene mutations result in both dyslipidemia and T2D, including *ABCA1* (Saleheen et al. 2006), *LIPE* (Albert *et al.* 2014), *LPL* (Hu *et al.* 2007), and *LRP6* (Mani *et al.* 2007). Genome-wide association studies (GWAS) have identified >150 loci associated with variation in plasma lipids (Teslovich *et al.* 2010),(Global Lipids Genetics Consortium *et al*. 2013) and >70 loci associated with T2D, fasting plasma glucose, glycated hemoglobin (HbA1c), or insulin resistance (Dupuis *et al.* 2010),(Soranzo *et al.* 2010),(Manning *et al.* 2012). Over a dozen of the loci detected are associated with both lipid and T2D-related traits at the genome-wide significance level, including APOB, GCKR, TIMD4, LPA, HLA-B, MLXIPL, NPC1L1, CYP7A1, FADS1–2-3, LRP1, LACTB, CETP, APOE-C1-C2, TOP1, and PLTP (http://www.genome.gov/GWAStudies/). Surprisingly, half of them, including CETP, MLXIPL, PLTP, GCKR, APOB, APOE-C1-C2, CYP7A1, and TIMD4, have shown opposite allelic effect on dyslipidemia and glucose levels (Li *et al.* 2014b), which is in contrary to the positive correlations observed at the clinical level. Furthermore, it is challenging to establish causality between genetic variants and complex traits in humans due to small gene effects, complex genetic structure, and environmental influences.

A complementary approach to finding genetic components in human disease is to use animal models. Apolipoprotein E-deficient (*Apoe*−/−) mice are a commonly used mouse model of dyslipidemia, with elevations in non-HDL cholesterol levels and reductions in HDL levels, even when fed a low fat chow diet (Shi *et al*. 2000),(Tian *et al*. 2005). High fat diet feeding aggravates dyslipidemia. We have found that *Apoe*−/− mice with certain genetic backgrounds develop significant hyperglycemia and T2D when fed a Western-type diet but become resistant with some other genetic backgrounds (Su *et al*. 2006a),(Li *et al*. 2011). BALB/cJ (BALB) and SM/J *Apoe*−/− mice exhibit differences in dyslipidemia and T2D-related phenotypes (Liu *et al*. 2015). In the present study, we performed quantitative trait locus (QTL) analysis using a female cohort derived from an intercross between BALB-*Apoe*−/− and SM-*Apoe*−/− mice to explore potential genetic connections between dyslipidemia and T2D.

## Methods

### Mice

BALB and SM *Apoe*-/- mice were created using the classic congenic breeding strategy, as previously described (Liu *et al*. 2015). BALB-*Apoe*-/- mice were crossed with SM-*Apoe*-/- mice to generate F1s, which were intercrossed by brother-sister mating to generate a female F2 cohort. Mice were weaned at 3 weeks of age onto a rodent chow diet. At 6 weeks of age, F2 mice were started on a Western diet containing 21% fat, 34.1% sucrose, 0.15% cholesterol, and 19.5% casein (Harlan Laboratories, TD 88137) and maintained on the diet for 12 weeks. All procedures were in accordance with current National Institutes of Health guidelines and approved by the institutional Animal Care and Use Committee.

### Measurements of plasma glucose and lipid levels

Mice were bled twice: once before initiation of the Western diet and once at the end of the 12-week feeding period. Mice were fasted overnight before blood was drawn from the retro-orbital plexus with the animals under isoflurane anesthesia. Plasma glucose was measured with a Sigma glucose (HK) assay kit according to the manufacturer′s instructions with a modification to include a longer incubation time. Briefly, 6 μl of plasma samples were incubated with 150 μl of assay reagent in a 96-well plate for 30 min at 30 ^°^C. The absorbance at 340 nm was read on a Molecular Devices (Menlo Park, CA) plate reader. The measurements of total cholesterol, HDL cholesterol, and triglyceride were performed as reported previously (Tian *et al*. 2005). Non-HDL cholesterol was calculated as the difference between total and HDL cholesterol.

### Genotyping

Genomic DNA was isolated from the tails of mice by using the phenol/chloroform extraction and ethanol precipitation method. The Illumina LD linkage panel consisting of 377 SNP loci was used to genotype 244 F2 mice. Microsatellite markers were typed for chromosome 8 where SNP markers were uninformative in distinguishing the parental origin of alleles. DNA samples from the two parental strains and their F1s served as controls. Uninformative SNPs were excluded from QTL analysis. SNP markers were also filtered based on the expected pattern in the control samples, and F2 mice were filtered based on 95% call rates in genotype calls. After filtration, 228 F2s and 144 markers were included in genome-wide QTL analysis.

### Statistical analysis

QTL analysis was performed using J/qtl and Map Manager QTX software as previously reported (Su *et al*. 2006a),(Su *et al*. 2006b),(Yuan *et al*. 2009). The expectation maximization (EM) algorithm was used to detect main effect loci for each trait. One thousand permutations of trait values were run to define the genome-wide LOD (logarithm of odds) score threshold needed for significant or suggestive linkage of each trait. Loci that exceeded the 95th percentile of the permutation distribution were defined as significant (*P*<0.05) and those exceeding the 37th percentile were suggestive (*P*<0.63).

### Prioritization of positional candidate genes

The Sanger SNP database (http://www.sanger.ac.uk/sanger/Mouse_SnpViewer/rel-1410) was used to prioritize candidate genes for overlapping QTLs affecting plasma glucose and HDL cholesterol levels on chromosome (Chr) 9, which were mapped in two or more crosses derived from different parental strains for either phenotype. We converted the original mapping positions in cM for the confidence interval to physical positions in Mb and then examined SNPs within the confidence interval. Probable candidate genes were defined as those with one or more SNPs in coding or upstream promoter regions that were shared by the parental strains carrying the “high” allele but were different from the parental strains carrying the “low” allele at a QTL, as previously reported (Rowlan *et al*. 2013b).

## Results

### Trait value distributions

Values of fasting plasma glucose, non-HDL cholesterol and triglyceride levels of F2 mice on both chow and Western diets and of HDL cholesterol level on the chow diet are normally or approximately normally distributed (Fig. 1). Values of square root-transformed HDL cholesterol levels on the Western diet show a normal distribution. These data were then analyzed to search for QTLs affecting the traits. Loci with a genome-wide suggestive or significant *P* value are presented in Table 1.

**Table 1.** Significant and suggestive QTLs for plasma glucose and lipid levels in female F_2_ mice derived from BALB-*Apoe*^-/-^ and SM-*Apoe*^-/-^ mice.

| Locus | Chr | Trait | LOD<sup>a</sup> | p-value<sup>b</sup> | Peak (cM) | 95% CI<sup>c</sup> | High allele | Mode of Inheritance<sup>d</sup> |
| --- | --- | --- | --- | --- | --- | --- | --- | --- |
| *Bglu16* | 9 | Glucose-C | 2.214 | 0.549 | 2.37 | 0.37-30.37 | B | Additive |
| *Bglu13* | 5 | Glucose-W | 2.1.8 | <0.63 | 67.4 | 45.4-80.03 | S | Recessive |
| - | 5 | Glucose-W | 3.198 | 0.097 | 101.24 | 29.40-101.24 | S | Additive |
| *Bglu16* | 9 | Glucose-W | 3.12 | <0.63 | 2.37 | 0-10.37 | B | Additive |
| ***Bglu17*** | 9 | Glucose-W | **5.425** | **0.001** | 26.37 | 16.37-40.37 | B | Additive |
| ***Hdlq5*** | 1 | HDL-C | **8.64** | **0.000** | 93.52 | 87.52-97.02 | S | Additive |
| *Hdlcl1* | 7 | HDL-C | 2.668 | 0.321 | 61.33 | 35.57-89.57 | S | Dominant |
| ***Hdlq17*** | 9 | HDL-C | **4.614** | **0.014** | 30.37 | 16.37-32.37 | S | Additive |
| ***Hdlq26*** | 10 | HDL-C | 2.181 | 0.591 | 61.22 | 25.03-61.22 | S | Dominant |
| ***Hdlq5*** | 1 | HDL-W | **13.944** | **0.000** | 87.52 | 83.52-93.52 | S | Additive |
| ***Hdlcl1*** | 7 | HDL-W | **3.658** | **0.034** | 85.57 | 77.57-89.67 | S | Additive |
| ***Hdlq17*** | 9 | HDL-W | **10.625** | **0.000** | 30.42 | 24.37-30.53 | S | Additive |
| *Chol7* | 1 | non-HDL-C | 2.093 | 0.626 | 66.95 | 9.52-74.56 | B | Recessive |
| *Nhdlq15* | 2 | non-HDL-C | 2.56 | 0.321 | 23.86 | 8.73-38.73 | B | Additive |
| *Hdlq34* | 5 | non-HDL-C | 2.106 | 0.614 | 19.4 | 19.4-30.5 | S | Additive |
| *Pnhdlc1* | 6 | non-HDL-C | 2.489 | 0.362 | 57.53 | 1.53-77.53 | B | Recessive |
| *Nhdlq1* | 8 | non-HDL-C | 2.221 | 0.537 | 44.14 | 10.14-60.14 | B | Additive |
| *Nhdlq12* | 12 | non-HDL-C | 2.73 | 0.245 | 39.41 | 15.41-59.41 | B | Additive |
| ***Nhdlq15*** | **2** | **non-HDL-W** | **4.79** | **0.002** | 31.80 | 22.73-40.73 | B | Dominant |
| *Nhdlq11* | 9 | non-HDL-W | 2.136 | 0.585 | 32.37 | 0.37-75.33 | B | Additive |
| - | 11 | non-HDL-W | 2.332 | 0.436 | 1.99 | 1.99-17.99 | B | Dominant |
| ***Nhdlq16*** | **16** | **non-HDL-W** | **3.99** | **0.011** | 46.66 | 35.43-46.66 | S | Dominant |
| *Tgql1* | 2 | Triglyceride-C | 2.952 | 0.169 | 26.73 | 12.73-60.83 | B | Additive |
| - | 5 | Triglyceride-C | 2.759 | 0.234 | 80.03 | 73.40-93.40 | S | Heterosis |
| *Trglyd* | 1 | Triglyceride-W | 3.291 | 0.091 | 97.02 | 79.24-97.02 | S | Additive |

<sup>a</sup> LOD scores were obtained from genome-wide QTL analysis using J/qtl software. The significant LOD scores are highlighted in bold. The suggestive and significant LOD score thresholds were determined by 1,000 permutation tests for each trait. Suggestive and significant LOD scores were 2.116 and 3.429, respectively, for glucose on the chow diet; 2.056 and 3.569 for glucose on the Western diet; 2.127 and 3.725 for HDL cholesterol, 2.09 and 3.662 for non-HDL cholesterol, and 2.102 and 3.522 for triglyceride on the chow diet; 2.10 and 3.486 for HDL, 2.123 and 3.628 for non-HDL, and 2.123 and 3.628 for triglyceride on the Western diet. <sup>b</sup> The p-values reported represent the level of genome-wide significance. <sup>c</sup> 95% Confidence interval in cM defined by a whole genome QTL scan. <sup>d</sup> Mode of inheritance was defined according to allelic effect at the nearest marker of a QTL. C: chow diet; W: Western diet.

**Figure 1.**
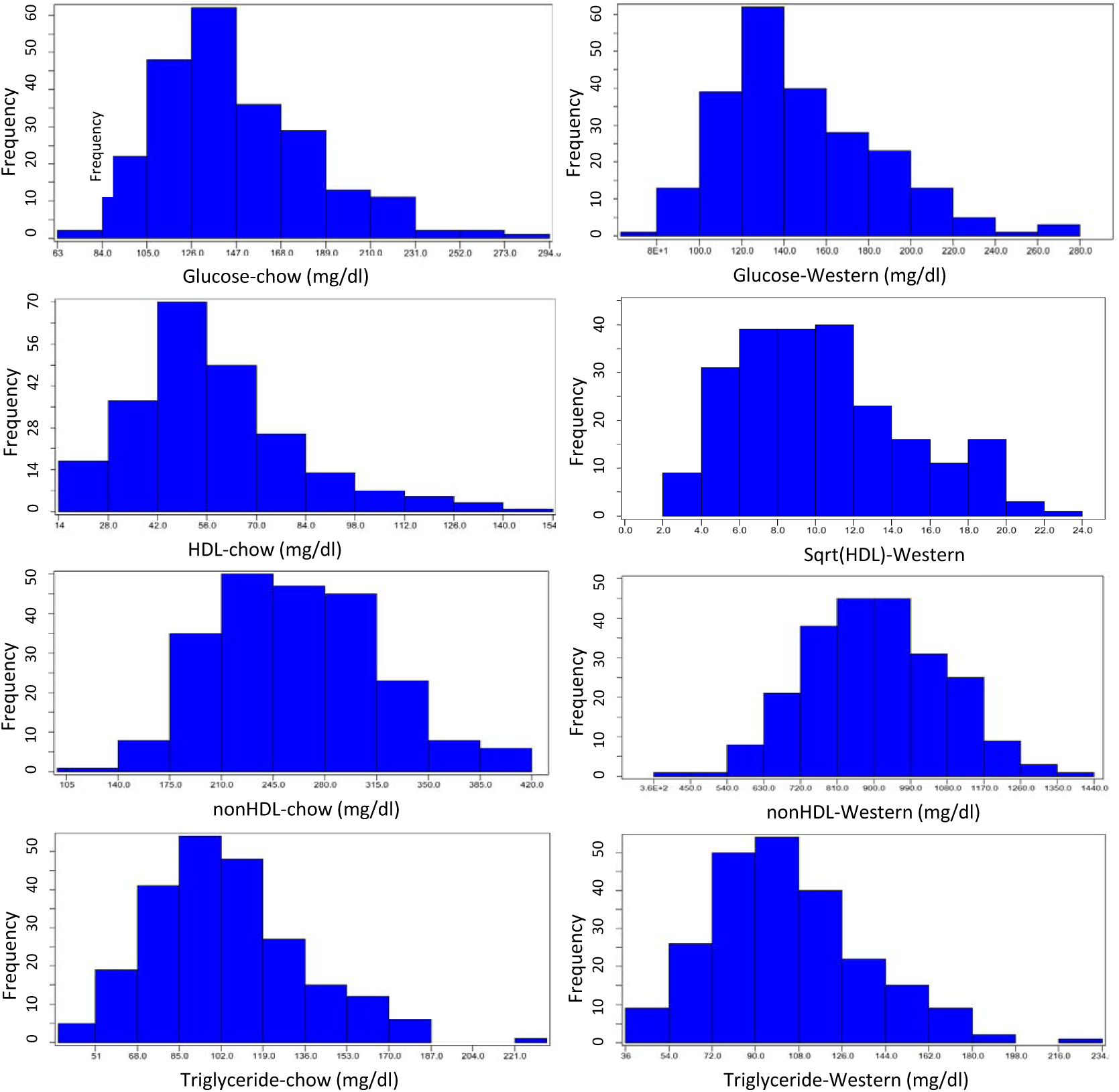
The distributions of trait values for fasting plasma glucose, HDL, non-HDL cholesterol and triglyceride of 228 female F2 mice derived from an intercross between BALB-*Apoe*^−/−^ and SM-*Apoe*^−/−^ mice. Fasting blood was collected once before initiation of the Western diet (left panel) and once after 12 weeks on the Western diet (right panel). Graphs were created using a plotting function of J/qtl software.

### Fasting glucose levels

A genome-wide scan for main effect QTL revealed a suggestive QTL near the proximal end of Chr9 for fasting glucose when mice were fed the chow diet (2.37 cM, LOD: 2.21) (Fig. 2 and Table 1). This QTL was replicated on the Western diet, and named *Bglu16.* For fasting glucose levels on the Western diet, a significant QTL on Chr9 and 3 suggestive QTLs, including *Bglu16* on Chr9, were identified. The significant QTL on Chr9 peaked at 26.37 cM and had a LOD score of 5.425. It was named *Bglu17.* The suggestive QTL near the middle portion of Chr5 (67.4 cM, LOD 2.18) replicated *Bglu13*, initially mapped in a B6 × BALB *Apoe*^-/-^ intercross (Zhang *et al*. 2012). The suggestive QTL on distal Chr5 (101.24 cM, LOD 3.198) was novel. The BALB allele conferred an increased glucose level for both of the Chr9 QTLs while the SM allele conferred increased glucose levels for the 2 Chr5 QTLs (Table 2).

**Table 2.** Allelic effects in different QTLs on plasma glucose and lipids of female F2 mice derived from BALB and SM *Apoe*^−/−^ mice.

| Locus name<sup>a</sup> | Chr | Trait | LOD | Peak (cM) | Closest marker | BB | SS | SB |
| --- | --- | --- | --- | --- | --- | --- | --- | --- |
| *Bglu16* | 9 | Glucose-C | 2.214 | 2.37 | rs13480073 | 109.0 ± 28.7 (n=44) | 93.9 ± 22.9 (n=43) | 97.4 ± 22.6 (n=141) |
| *Bglu13* | 5 | Glucose-W | 2.1.8 | 67.4 | rs3726547 | 144.4 ± 30.6 (n=43) | 153.5 ± 40.4 (n=88) | 142.9 ± 35.3 (n=97) |
| - | 5 | Glucose-W | 3.198 | 101.24 | rs13478578 | 132.7 ± 31.3 (n=51) | 158.8 ± 41.8 (n=63) | 147.5 ± 34.0 (n=113) |
| *Bglu16* | 9 | Glucose-W | 3.12 | 2.37 | rs13480073 | 165.3 ± 40.9 (n=54) | 138.3 ± 30.0 (n=43) | 144.4 ± 35.7 (n=141) |
| ***Bglu17*** | 9 | Glucose-W | **5.425** | 26.37 | CEL.9_49183636 | 168.0 ± 39.8 (n=42) | 134.7 ± 26.5 (n=62) | 146.1 ± 37.1 (n=124) |
| ***Hdlq5*** | 1 | HDL-C | **8.64** | 93.52 | rs13476259 | 49.5 ± 20.9 (n=60) | 73.2 ± 26.5 (n=62) | 55.1 ± 19.4 (n=106) |
| *Hdlcl1* | 7 | HDL-C | 2.668 | 61.33 | rs3724711 | 49.0 ± 20.0 (n=50) | 58.3 ± 21.0 (n=63) | 62.9 ± 25.5 (n=115) |
| ***Hdlq17*** | 9 | HDL-C | **4.614** | 30.37 | CEL.9_49183636 | 49.5 ± 15.9 (n=42) | 69.4 ± 26.8 (n=62) | 56.2 ± 22.5 (n=124) |
| *Hdlq26* | 10 | HDL-C | 2.181 | 61.22 | rs3688351 | 50.7 ± 19.2 (n=60) | 59.4 ± 21.9 (n=53) | 62.4 ± 25.8 (n=114) |
| ***Hdlq5*** | 1 | sqrtHDL-W | **13.944** | 87.52 | r(Wang *et al.* 2004)s3685643 | 66.6 ± 50.2 (n=57) | 201.1 ± 118.8 (n=62) | 117.0 ± 97.1 (n=109) |
| ***Hdlcl1*** | 7 | sqrtHDL-W | **3.658** | 85.57 | rs6216320 | 95.5 ± 87.6 (n=63) | 173.3 ± 126.7 (n=55) | 122.4 ± 98.4 (n=110) |
| ***Hdlq17*** | 9 | sqrtHDL-W | **10.625** | 30.42 | CEL.9_49183636 | 57.6 ± 49.3 (n=42) | 183.3 ± 115.0 (n=62) | 122.8 ± 101.7 (n=124) |
| *Chol7* | 1 | non-HDL-C | 2.093 | 66.95 | rs6354736 | 279.5 ± 62.8 (n=56) | 257.8 ± 56.9 (n=57) | 251.2 ± 52.1 (n=114) |
| *Nhdlq15* | 2 | non-HDL-C | 2.56 | 23.86 | mCV23209429 | 273.9 ± 56.1 (n=55) | 238.0 ± 47.0 (n=53) | 262.7 ± 59.0 (n=120) |
| *Hdlq34* | 5 | non-HDL-C | 2.106 | 19.4 | rs3658401 | 244.5 ± 54.7 (n=63) | 276.0 ± 53.2 (n=61) | 259.3 ± 58.3 (n=104) |
| *Pnhdlc1* | 6 | non-HDL-C | 2.489 | 57.53 | rs13478909 | 279.6 ± 51.3 (n=51) | 252.0 ± 65.2 (n=57) | 254.8 ± 53.5 (n=120) |
| *Nhdlq1* | 8 | non-HDL-C | 2.221 | 44.14 | D8Mit50 | 275.0 ± 54.5 (n=60) | 242.4 ± 57.4 (n=57) | 262.9 ± 55.6 (n=96) |
| *Nhdlq12* | 12 | non-HDL-C | 2.73 | 39.41 | rs6195664 | 278.6 ± 52.3 (n=62) | 243.8 ± 57.8 (n=59) | 257.4 ± 56.5 (n=107) |
| ***Nhdlq15*** | **2** | non-HDL-W | **4.79** | 31.8 | rs13476507 | 954.1 ± 156.0 (n=56) | 806.9 ± 158.2 (n=47) | 915.6 ± 166.1 (n=125) |
| *Nhdlq11* | 9 | non-HDL-W | 2.136 | 32.37 | rs3709825 | 958.4 ± 211.4 (n=42) | 856.8 ± 165.3 (n=62) | 906.6 ± 149.5 (n=124) |
| - | 11 | non-HDL-W | 2.332 | 1.99 | rs4222040 | 927.3 ± 149.8 (n=67) | 849.0 ± 165.1 (n=69) | 917.0 ± 170.5 (n=85) |
| ***Nhdlq16*** | **16** | non-HDL-W | **3.99** | 46.66 | rs3721202 | 820.2 ± 152.7 (n=56) | 931.9 ± 146.4 (n=52) | 928.4 ± 174.6 (n=120) |
| *Tgql1* | 2 | Triglyceride-C | 2.952 | 26.73 | mCV23209429 | 123.7 ± 35.7 (n=55) | 101.9 ± 34.6 (n=53) | 107.3 ± 31.8 (n=120) |
| - | 5 | Triglyceride-C | 2.759 | 80.03 | gnf05.120.578 | 110.2 ± 33.2 (n=43) | 119.3 ± 35.8 (n=88) | 101.6 ± 31.3 (n=97) |
| *Trglyd* | 1 | Triglyceride-W | 3.291 | 97.02 | rs13476259 | 94.0 ± 28.6 (n=59) | 115.3 ± 33.0 (n=62) | 100.7 ± 30.8 (n=106) |

Data are mean ± SD. The units for these measurements are mg/dL for plasma glucose or lipid levels. The number in the brackets represents the number of progeny with a specific genotype at a peak marker. The significant QTLs and their LOD scores were highlighted in bold. Chr, chromosome; LOD, logarithm of odds; C, chow diet; W, Western diet; BB, homozygous BALB allele; SS, homozygous SM allele; SM, heterozygous allele.

**Figure 2.**
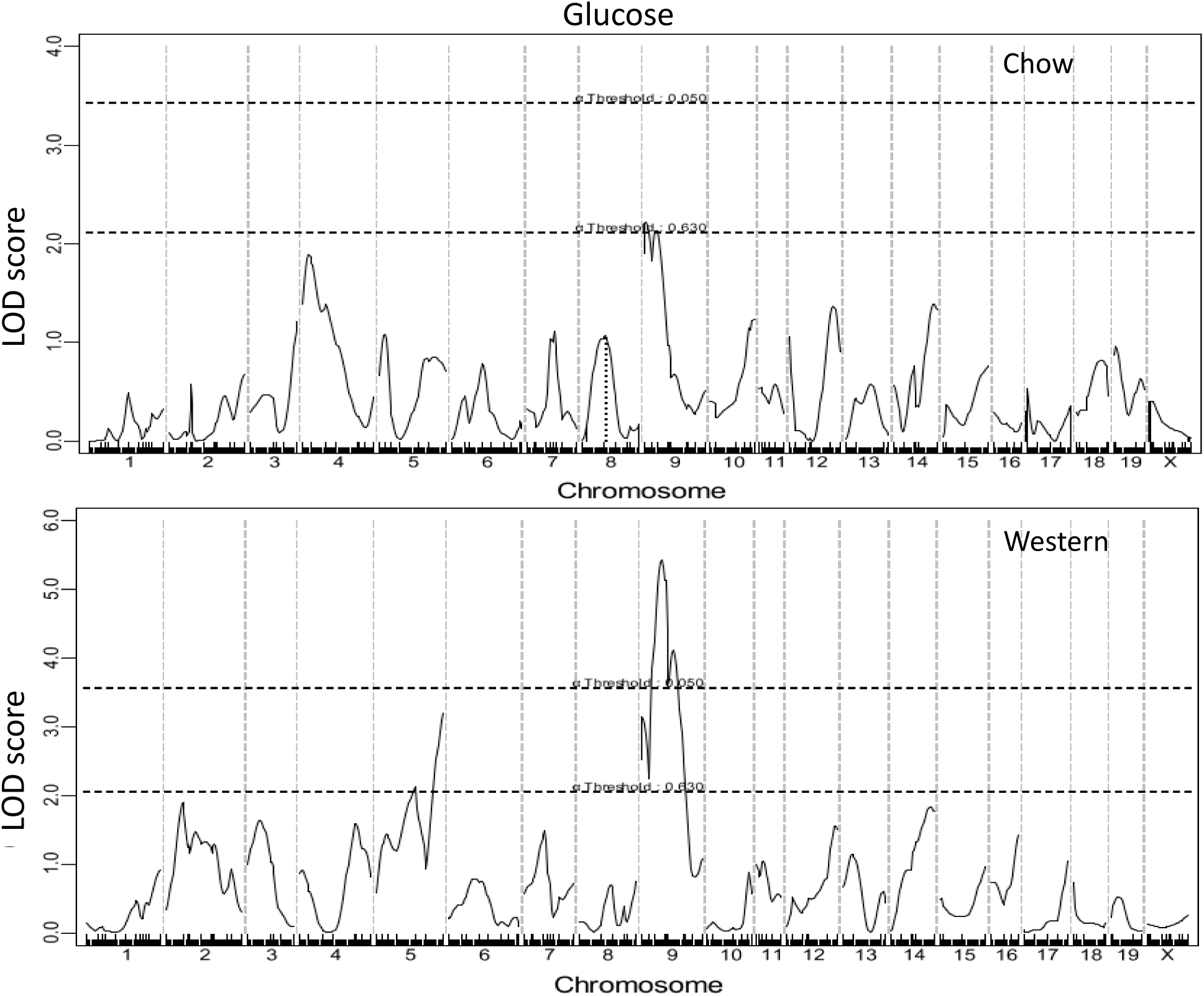
Genome-wide scans to search for main effect loci influencing fasting plasma glucose levels of female F2 mice when fed a chow or Western diet. Chromosomes 1 through 20 are represented numerically on the X-axis. The Y-axis represents the LOD score. Two horizontal dashed lines denote genome-wide empirical thresholds for suggestive (*P*=0.63) and significant (*P*=0.05) linkage.

### Fasting lipid levels

Genome-wide scans for main effect QTLs showed that HDL, non-HDL cholesterol, and triglyceride levels were each controlled by multiple QTLs (Fig. 3 through 5, Table 1). For HDL, 3 significant QTLs, located on Chr1, Chr7 and Chr9, and 1 suggestive QTL on Chr10, were identified. All 3 significant QTLs for HDL were detected when mice were fed either chow or Western diet, while the suggestive QTL on Chr10 was found when mice were fed the chow diet. The significant QTL on Chr1 replicated *Hdlq5,* which had been mapped in numerous crosses (Wang and Paigen. 2005). The Chr7 QTL replicated *Hdlcl1,* initially mapped in (PERA/EiJ × B6-*Ldlr*) × B6-*Ldlr* backcross (Seidelmann *et al*. 2005). The Chr9 QTL replicated *Hdlq17*, previously mapped in B6 × 129S1/SvImJ F2 mice (Ishimori *et al*. 2004b). The suggestive QTL on Chr10 overlapped with *Hdlq26* mapped in a SM/J × NZB/BlNJ intercross (Korstanje *et al*. 2004). For all 4 HDL QTLs, F2 mice homozygous for the SS allele had higher HDL levels than those homozygous for the BB allele (Table 2).

**Figure 3.**
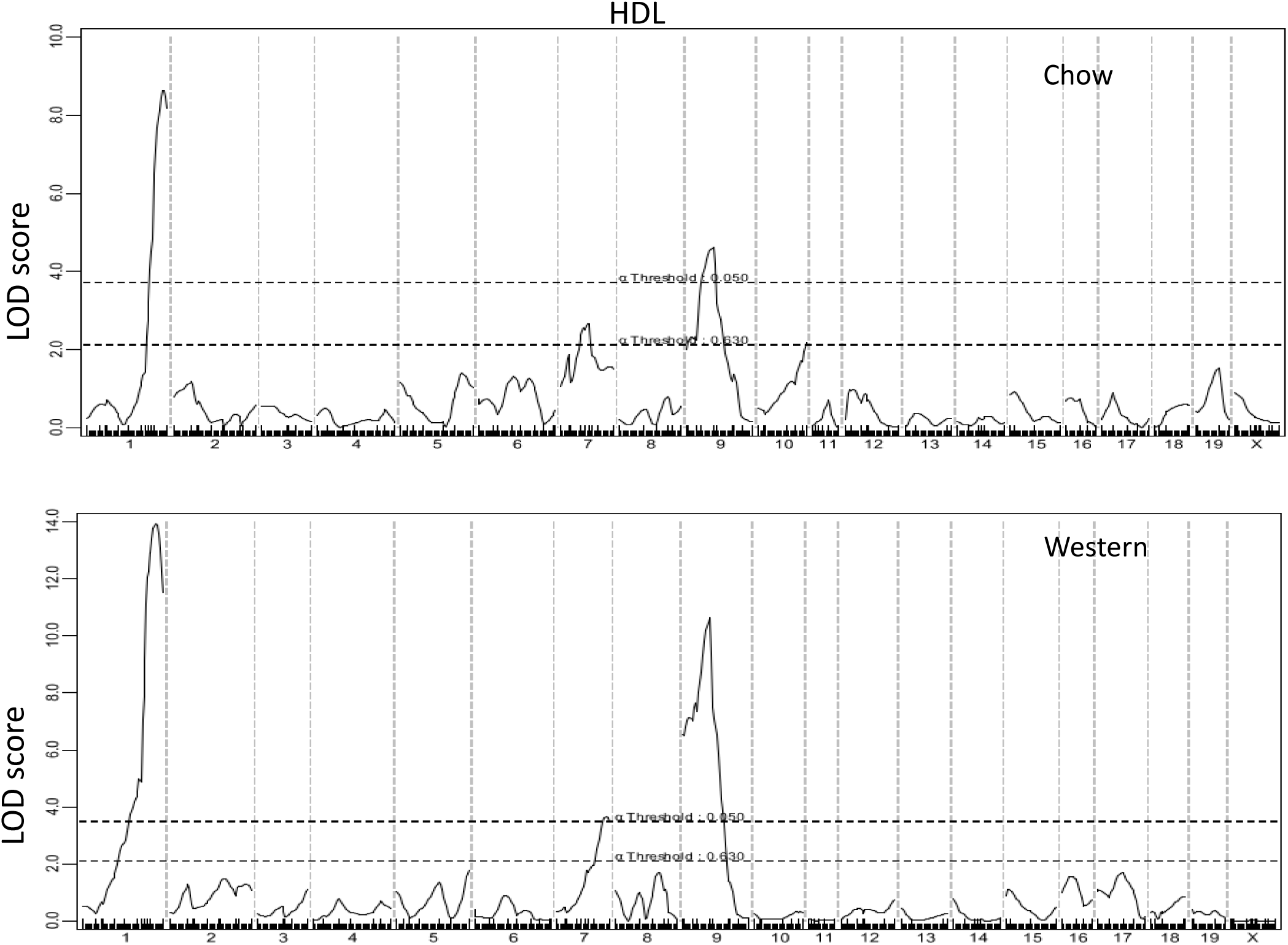
Genome-wide scans to search for loci influencing HDL cholesterol levels of female F2 mice when fed a chow or Western diet. Three significant loci on chromosomes 1, 7, and 9 and one suggestive locus on chromosome 10 were detected to affect HDL cholesterol levels of mice.

For non-HDL cholesterol levels, 6 suggestive QTLs were detected when F2 mice were fed the chow diet, and 2 significant and 2 suggestive QTLs were detected on the Western diet (Fig. 4). The 2 significant QTLs on Chr2 and Chr16 and the suggestive QTL on Chr11 were novel. The significant QTLs on Chr2 and Chr16 were named *Nhdlq15* and *Nhdlq16*, respectively. *Nhdlq15* peaked at 31.8 cM on Chr2 and affected non-HDL levels in a dominant mode from the BB allele while *Nhdlq16* peaked at 46.66 cM on Chr16 and affected non-HDL levels in a dominant mode from the SS allele. The rest replicated previously identified ones in other mouse crosses: The Chr1 QTL peaked at 66.95 cM, overlapping with *Chol7* mapped in an intercross of 129S1/SvImJ and CAST/Ei mice (Lyons *et al*. 2004). The Chr5 QTL overlapped with *Hdlq34* mapped in PERA/EiJ × I/LnJ and PERA/EiJ × DBA/2J intercrosses (Wittenburg *et al*. 2006). The Chr6 QTL overlapped with *Pnhdlc1*, initially mapped in a B6 × CASA/Rk intercross and then replicated in B6 × C3H *Apoe*^-/-^ F2 mice (Sehayek *et al*. 2003),(Li *et al*. 2008). The Chr8 QTL replicated *Nhdlq1*, initially mapped in B6 × 129S1/SvImJ F2 mice (Ishimori *et al*. 2004a). The Chr9 QTL replicated *Nhdlq11*, initially mapped in B6 × C3H *Apoe*^-/-^ F2 mice (Li *et al*. 2008). The Chr12 QTL peaked at 44.14 cM, overlapping with *Nhdlq12* mapped in a B6 × C3H *Apoe*^-/-^ F2 intercross (Li *et al*. 2008).

**Figure 4.**
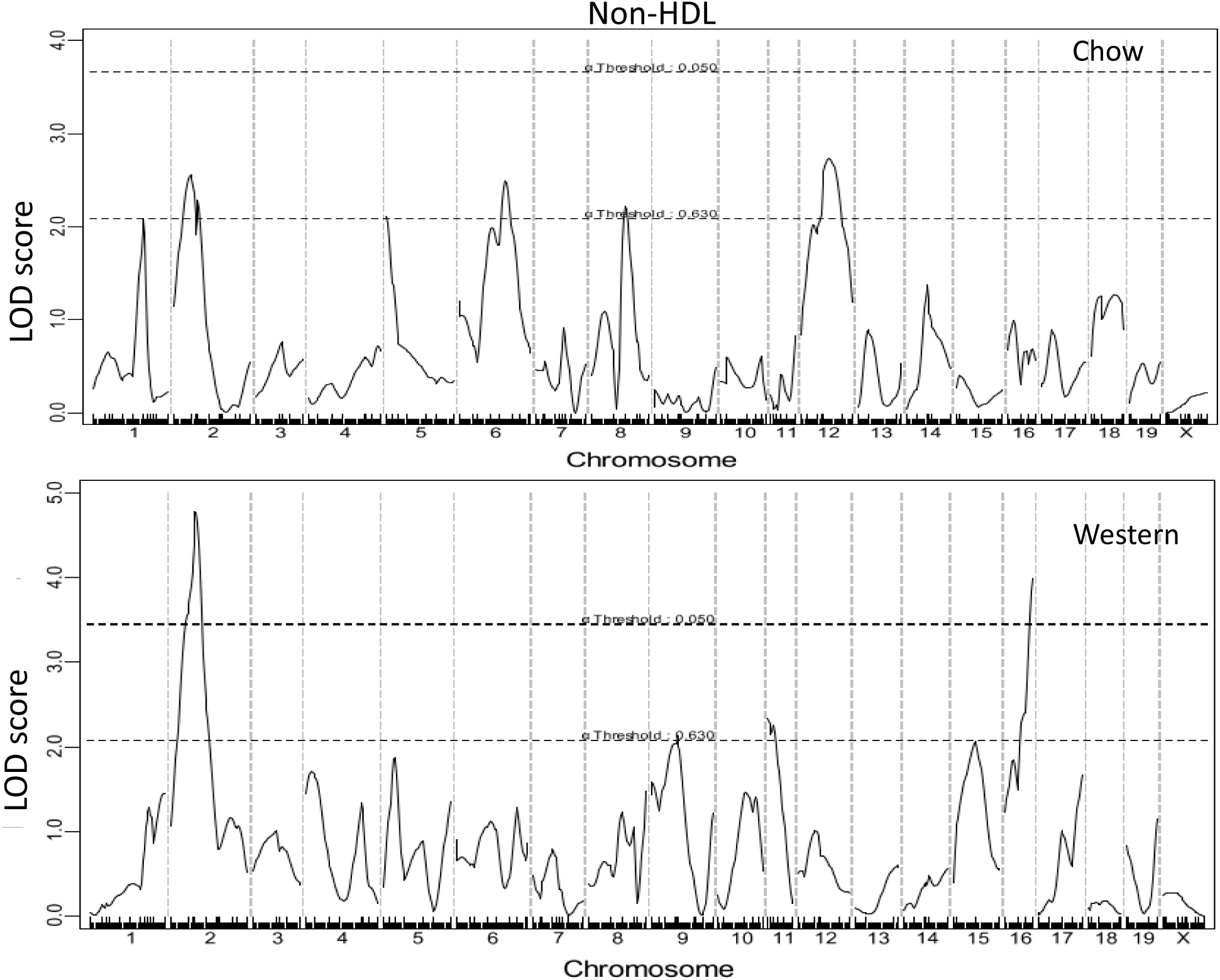
Genome-wide scans to search for loci influencing non-HDL cholesterol levels of female F2 mice fed a chow or Western diet. Two significant loci on chromosomes 2 and 16 were identified to affect non-HDL cholesterol levels of mice fed the Western diet.

For triglyceride levels, 3 suggestive QTLs, located on Chr1, 2, and 5, respectively, were identified (Fig. 5). The Chr1 QTL peaked at 97 cM, 17 cM distal the *Apoa2* gene (80 cM). The Chr2 QTL replicated *Tgq11,* mapped in an intercross between DBA/1J and DBA/2J (Stylianou *et al*. 2008). The Chr5 QTL was novel.

**Figure 5.**
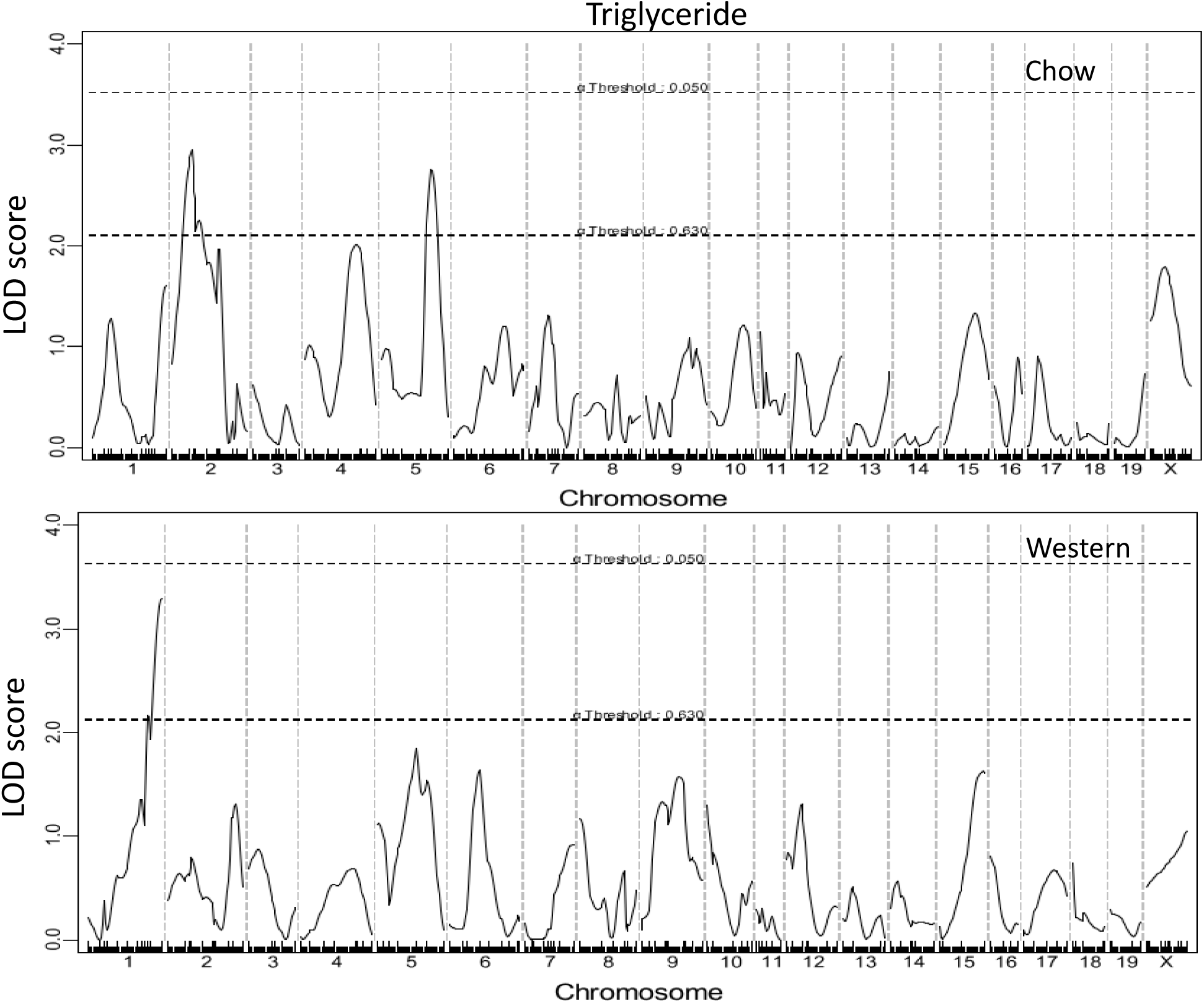
Genome-wide scans to search for loci influencing triglyceride levels of female F2 mice fed a chow or Western diet. Three suggestive loci were identified for triglyceride levels.

### Coincident QTLs for fasting glucose and lipids

LOD score plots for Chr9 showed that the QTL for fasting glucose (*Bglu17*) coincided precisely with the QTLs for HDL (*Hdlq17*) and non-HDL (*Nhdlq11*) in the confidence interval (Fig. 6). F2 mice homozygous for the BB allele exhibited elevated levels of fasting glucose and non-HDL but decreased levels of HDL, compared to those homozygous for the SS allele (Table 2). These QTLs affected their respective trait values in an additive manner.

**Figure 6.**
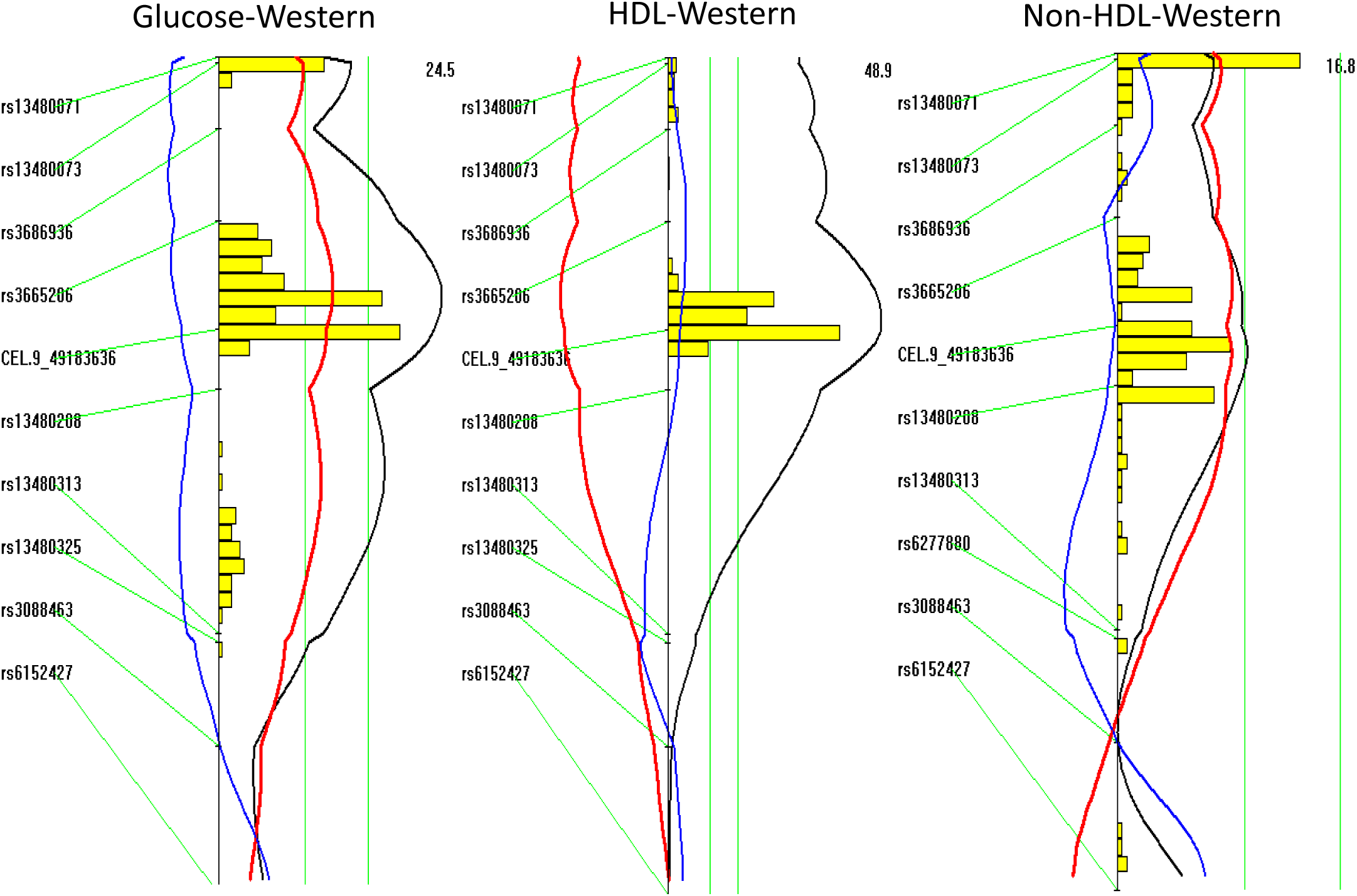
LOD score plots for fasting glucose, HDL, and non-HDL cholesterol of F2 mice fed the Western diet on chromosome 9. Plots were created with the interval mapping function of Map Manager QTX. The histogram in the plot estimates the confidence interval for a QTL. Two green horizontal lines represent genome-wide significance thresholds for suggestive or significant linkage (*P*=0.63 and *P*=0.05, respectively). Black plots reflect the LOD score calculated at 1-cM intervals, the red plot represents the effect of the BALB allele, and the blue plot represents the effect of the SM allele. If BALB represents the high allele, then the red plot will be to the right of the graph; otherwise, it will be to the left.

### Correlations between plasma glucose and lipid levels

The correlations of fasting glucose levels with plasma levels of HDL, non-HDL cholesterol, or triglyceride were analyzed with the F2 population (Fig. 7). A significant inverse correlation between fasting glucose and HDL cholesterol levels was observed when the mice were fed a chow (*R*=-0.220; *P*=8.1E-4) or Western diet (*R*=-0.257; *P*=8.5E-5). F2 mice with higher HDL cholesterol levels had lower fasting glucose levels. Conversely, significant positive correlations between fasting glucose and non-HDL cholesterol levels were observed when mice were fed either chow (*R*=0.194; *P*=3.31E-3) or Western diet (*R*=0.558; *P*=4.7E-20). F2 mice with higher non-HDL cholesterol levels also had higher fasting glucose levels, especially on the Western diet. A significant positive correlation between plasma levels of fasting glucose and triglyceride was observed when mice were fed the Western diet (*R*=0.377; *P*=3.9E-9) but not the chow diet (*R*=0.065; *P*=0.330).

### Prioritization of positional candidate genes for Chr9 coincident QTLs

*Bglu17* on Chr9 has been mapped in 3 separate intercrosses, including previously reported C57BLKS × DBA/2 (Yaguchi *et al*. 2005) and B6-*Apoe*^-/-^ × BALB-*Apoe*^-/-^ crosses (Zhang *et al*. 2012). *Hdlq17* on Chr9 has been mapped in multiple crosses, including B6 × 129, B6 × CAST/EiJ, 129 × RIII, B6-*Apoe*^-/-^ × C3H-*Apoe*^-/-^, and B6-*Apoe*^-/-^ × BALB-*Apoe*^-/-^ crosses (Sehayek *et al*. 2003),(Su *et al*. 2009),(Lyons *et al*. 2004)(Lyons *et al*. 2003)(Rowlan *et al*. 2013b)(Rowlan *et al*. 2013a). The latter 2 crosses were not included in haplotype analysis because the BALB or C3H allele at the locus is associated with elevated HDL cholesterol levels, opposite to the current observation. We conducted haplotype analyses using Sanger SNP database to prioritize positional candidate genes for both QTLs. Prioritized candidate genes for *Hdlq17* are shown in Supplementary Table 1, and candidate genes for *Bglu17* are shown in Supplementary Table 2. Some candidates for *Hdlq17* are also candidate genes for *Bglu17*, including *Phldb1*, *Mill1, Ube4a, Cd3e, Sik3, Apoc3, Dixdc1*, and *Alg9*. Potential candidate genes contain one or more non-synonymous SNPs in the coding regions or SNPs in the upstream regulatory region that are shared by the high allele strains but are different from the low allele strains at the QTL. All candidate genes were further examined for associations with relevant human diseases using the NIH GWAS database (http://www.genome.gov/GWAStudies/) (Global Lipids Genetics Consortium *et al*. 2013). *Phldb1, Sik3,* and *Apoc3* have been shown to be associated with variations in total, HDL, LDL-cholesterol or triglyceride levels (Global Lipids Genetics Consortium *et al*. 2013),(Ko *et al*. 2014),(Teslovich *et al*. 2010).

**Figure 7.**
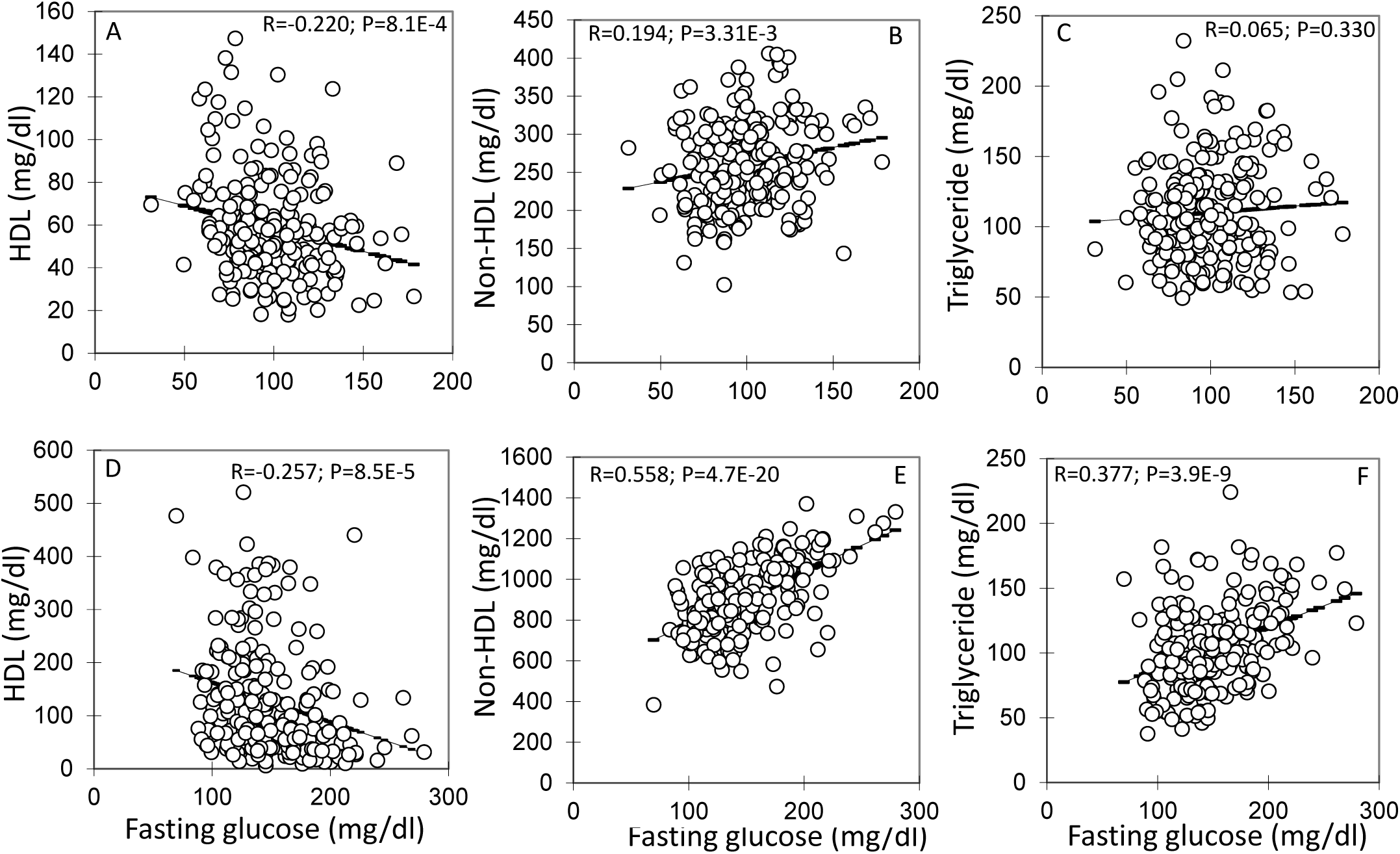
Correlations of fasting plasma glucose levels with plasma levels of HDL, non-HDL cholesterol and triglyceride in the F2 population fed a chow (A, B and C) or Western diet (D, E and F). Each point represents values of an individual F_2_ mouse. The correlation coefficient (*R*) and significance (*P*) are shown.

## Discussion

BALB and SM are two mouse strains that exhibit distinct differences in HDL, non-HDL cholesterol, and type 2 diabetes-related traits when deficient in *Apoe* (Liu *et al*. 2015). BALB-*Apoe*^-/-^ mice have higher HDL, lower non-HDL cholesterol, and lower glucose levels than SM-*Apoe*^-/-^ mice when fed a Western diet. To identify the genetic factors underlying these differences, we performed QTL analysis on a female cohort derived from an intercross between the two *Apoe*^-/-^ strains. We have identified four loci for fasting glucose levels, four loci for HDL cholesterol levels, nine loci for non-HDL cholesterol levels, and three loci for triglyceride levels. Moreover, we have observed the colocalization of QTL for fasting glucose with the QTLs for HDL and non-HDL cholesterol on chromosome 9 and the sharing of potential candidate genes.

We identified a significant QTL near 26 cM on chromosome 9, which affected fasting plasma glucose levels of mice on either chow or Western diet. We named it *Bglu17* to represent a novel locus regulating fasting glucose levels in the mouse. This locus is overlapping with a significant QTL (not named) for blood glucose levels on the intraperitoneal glucose tolerance test identified in a BKS.Cg-*Leprdb*+/+*m ×* DBA/2 intercross and a suggestive QTL identified in a B6-*Apoe*^-/-^ × BALB-*Apoe*^-/-^ intercross (Zhang *et al*. 2012),(Yaguchi *et al*. 2005). Interestingly, we found that *Bglu17* coincided precisely with *Hdlq17*, a QTL for HDL cholesterol levels, and *Nhdlq11*, a QTL for non-HDL cholesterol levels. The colocalization of two or more QTLs for different traits suggests that these traits are controlled either by the same gene(s) or closely linked but different individual genes. *Hdlq17* has been mapped in multiple crosses derived from inbred mouse strains whose genomes have been resequenced by Sanger, including B6, 129, BALB, and CAST/EiJ (Rowlan *et al*. 2013b),(Ishimori *et al*. 2004b),(Sehayek *et al*. 2003),(Li *et al*. 2008),(Lyons *et al*. 2004),(Su *et al*. 2009),(Rowlan *et al*. 2013a). *Nhdlq11* was previously mapped in DBA/2 × CAST/EiJ, NZB/BIN × SM, and B6-*Apoe*^-/-^ × C3H-*Apoe*^-/-^ intercrosses (Li *et al*. 2008),(Purcell-Huynh *et al*. 1995)(Lyons *et al*. 2003). To determine whether *Bglu17* and *Hdlq17* share the same underlying candidate genes, we performed haplotype analyses on those crosses that led to the identification of the QTLs. Eight candidate genes are shared by both QTLs. Of them, *Sik3* and *Apoc3* are located precisely underneath the linkage peak of *Bglu17* and *Hdlq17*, and they are also functional candidate genes of *Hdlq17*. Indeed, recent GWAS studies have associated these 2 genes with dyslipidemia or variations in HDL, LDL cholesterol, and triglyceride levels (Teslovich *et al*. 2010),(Ko *et al*. 2014),(Willer and Mohlke. 2012). These “lipid genes” might also be the causal gene(s) of *Bglu17*, contributing to variation in fasting glucose levels. Although it is unknown how they affect glucose homeostasis, one probable pathway is through their action on plasma lipid levels, which then predispose variation in glucose-related traits. The current observation on the significant correlations of HDL, non-HDL cholesterol, and triglyceride levels with fasting glucose levels in this cross supports this speculation. Plasma lipid levels, especially non-HDL cholesterol, of the F2 mice were significantly elevated on the Western diet, so were the fasting glucose levels. When fed the Western diet, *Apoe*^-/-^ mice display a rapid rise in non-HDL cholesterol levels, often reaching a plateau within a couple of weeks (unpublished data), whereas their blood glucose levels rise more slowly and gradually within 12 weeks (Li *et al*. 2012),(Zhou *et al*. 2015). The early onset suggests that hyperlipidemia may play a causal role in the rise of blood glucose in the *Apoe*^-/-^ mouse model.

A significant reverse correlation was observed between plasma HDL cholesterol levels and fasting glucose levels in this cross on either chow or Western diet. This result is consistent with the findings of prospective human studies that low HDL levels can predict the future risk of developing T2D and low HDL levels are more prevalent in diabetic patients than in the normal population (Wilson *et al*. 1985),(Wilson *et al*. 2007). HDL can increase insulin secretion from β-cells, improve insulin sensitivity of the target tissues, and accelerate glucose uptake by muscle via the AMP-activated protein kinase (Drew *et al*. 2012). A significant correlation of non-HDL cholesterol levels with fasting glucose levels was also observed in this cross, and the significance of correlation was extremely high when mice were fed the Western diet. Emerging human studies have also revealed associations of non-HDL cholesterol and ApoB with fasting glucose levels and incident type 2 diabetes (Hwang *et al*. 2014b),(Hwang *et al*. 2014a),(Ley *et al*. 2010). We previously observed that the elevation of non-HDL cholesterol levels in *Apoe*^-/-^ mice during consumption of a Western diet induces a chronic, low-grade inflammation state characterized by rises in circulating cytokines and infiltration of monocytes/macrophages in various organs or tissues (Tian *et al*. 2005),(Li *et al*. 2011),(Li *et al*. 2012)(Zhang *et al*. 2015). Inflammation in the islets impairs β-cell function (Li *et al*. 2011). LDL can also directly affect function and survival of β-cells (Rutti *et al*. 2009). In addition, high levels of LDL can induce insulin resistance due to its lipotoxicity and effect on endoplasmic reticulum stress (Li *et al*. 2014a).

Plasma triglyceride levels were strongly correlated with fasting glucose levels in this cross on the Western diet, although no significant correlation was found when mice were fed the chow diet. Despite the strong correlation, no overlapping QTLs were found for fasting glucose and triglyceride. The reason for the discrepancy between non-HDL cholesterol and triglyceride in terms of the presence or absence of colocalized QTLs is unclear.

A suggestive QTL for fasting glucose near the proximal end of chromosome 9 (2.37 cM) was detected in this cross, initially on the chow diet and then replicated on the Western diet. The LOD score plot for chromosome 9 has shown 2 distinct peaks, one with a suggestive LOD score at the proximal end and one with a significant LOD score at a more distal region, suggesting the existence of two loci for fasting glucose on the chromosome. The bootstrap test, a statistical method for defining the confidence interval of QTLs using simulation (Visscher *et al*. 1996), also indicated the existence of two QTLs for the trait on chromosome 9. We named the proximal one *Bglu16* to represent a new QTL for fasting glucose in the mouse. Naming a suggestive locus is considered appropriate if it is repeatedly observed (Abiola *et al*. 2003).

Two suggestive QTLs for fasting glucose on chromosome 5 were identified when mice were fed the Western diet. The proximal one replicated *Bglu13*, recently mapped in the B6-*Apoe*^-/-^ × BALB-*Apoe*^-/-^ cross (Zhang *et al*. 2012). One probable candidate gene for this QTL is *Hnf1a*, which encodes hepatocyte nuclear factor 1α. In humans, *Hnf1a* mutations are the most common cause of maturity-onset diabetes of the young (MODY) (Shepherd *et al*. 2001). The suggestive QTL in the distal region was novel.

Most of the QTLs identified for plasma lipids confirm those identified in previous studies, whereas two QTLs for non-HDL are new and named *Nhdlq15* and *Nhdlq16*. The QTLs on distal chromosome 1 for HDL and triglyceride has been mapped in a number of mouse crosses, and *Apoa2* has been identified as the underlying causal gene (Wang *et al*. 2004). However, the QTL (∼90 cM) mapped in this study showed that it was more distal to the *Apoa2* gene (80 cM), thus suggesting a different underlying causal gene.

In summary, we have identified multiple QTLs contributing to dyslipidemia and hyperglycemia in a segregating F2 population. The findings on the colocalization of QTLs for fasting glucose, HDL and non-HDL cholesterol levels and the sharing of potential candidate genes demonstrate genetic connections between dyslipidemia and type 2 diabetes. The significant correlations of fasting glucose with HDL, non-HDL cholesterol, and triglyceride observed in this study support the hypothesis that dyslipidemia plays a causal role in the development of type 2 diabetes (Li *et al*. 2014a).

## LITERATURE CITED

Abiola, O., J. M. Angel, P. Avner, A. A. Bachmanov, J. K. Belknap et al, 2003 The nature and identification of quantitative trait loci: A community’s view. Nat. Rev. Genet. 4: 911–916.

Albert, J. S., L. M. Yerges-Armstrong, R. B. Horenstein, T. I. Pollin, U. T. Sreenivasan et al, 2014 Null mutation in hormone-sensitive lipase gene and risk of type 2 diabetes. N. Engl. J. Med. 370: 2307–2315.

Drew, B. G., K. A. Rye, S. J. Duffy, P. Barter and B. A. Kingwell, 2012 The emerging role of HDL in glucose metabolism. Nat. Rev. Endocrinol. 8: 237–245.

Dupuis, J., C. Langenberg, I. Prokopenko, R. Saxena, N. Soranzo et al, 2010 New genetic loci implicated in fasting glucose homeostasis and their impact on type 2 diabetes risk. Nat. Genet. 42: 105–116.

Global Lipids Genetics Consortium, C. J. Willer, E. M. Schmidt, S. Sengupta, G. M. Peloso et al, 2013 Discovery and refinement of loci associated with lipid levels. Nat. Genet. 45: 1274–1283.

Hu, Y., Y. Ren, R. Z. Luo, X. Mao, X. Li et al, 2007 Novel mutations of the lipoprotein lipase gene associated with hypertriglyceridemia in members of type 2 diabetic pedigrees. J. Lipid Res. 48: 1681–1688.

Hwang, Y. C., H. Y. Ahn, S. W. Park and C. Y. Park, 2014a Apolipoprotein B and non-HDL cholesterol are more powerful predictors for incident type 2 diabetes than fasting glucose or glycated hemoglobin in subjects with normal glucose tolerance: A 3.3-year retrospective longitudinal study. Acta Diabetol. 51: 941–946.

Hwang, Y. C., H. Y. Ahn, S. H. Yu, S. W. Park and C. Y. Park, 2014b Atherogenic dyslipidaemic profiles associated with the development of type 2 diabetes: A 3.1-year longitudinal study. Diabet. Med. 31: 24–30.

Ishimori, N., R. Li, P. M. Kelmenson, R. Korstanje, K. A. Walsh et al, 2004a Quantitative trait loci that determine plasma lipids and obesity in C57BL/6J and 129S1/SvImJ inbred mice. J. Lipid Res. 45: 1624–1632.

Ishimori, N., R. Li, P. M. Kelmenson, R. Korstanje, K. A. Walsh et al, 2004b Quantitative trait loci analysis for plasma HDL-cholesterol concentrations and atherosclerosis susceptibility between inbred mouse strains C57BL/6J and 129S1/SvImJ. Arterioscler. Thromb. Vasc. Biol. 24: 161–166.

Ko, A., R. M. Cantor, D. Weissglas-Volkov, E. Nikkola, P. M. Reddy et al, 2014 Amerindian-specific regions under positive selection harbour new lipid variants in latinos. Nat. Commun. 5: 3983.

Korstanje, R., R. Li, T. Howard, P. Kelmenson, J. Marshall et al, 2004 Influence of sex and diet on quantitative trait loci for HDL cholesterol levels in an SM/J by NZB/BlNJ intercross population. J. Lipid Res. 45: 881–888.

Ley, S. H., S. B. Harris, P. W. Connelly, M. Mamakeesick, J. Gittelsohn et al, 2010 Association of apolipoprotein B with incident type 2 diabetes in an aboriginal canadian population. Clin. Chem. 56: 666–670.

Li, J., Z. Lu, Q. Wang, Z. Su, Y. Bao et al, 2012 Characterization of Bglu3, a mouse fasting glucose locus, and identification of apcs as an underlying candidate gene. Physiol. Genomics 44: 345–351.

Li, J., Q. Wang, W. Chai, M. H. Chen, Z. Liu et al, 2011 Hyperglycemia in apolipoprotein E-deficient mouse strains with different atherosclerosis susceptibility. Cardiovasc. Diabetol. 10: 117.

Li, N., J. Fu, D. P. Koonen, J. A. Kuivenhoven, H. Snieder et al, 2014a Are hypertriglyceridemia and low HDL causal factors in the development of insulin resistance? Atherosclerosis 233: 130–138.

Li, N., M. R. van der Sijde, LifeLines Cohort Study Group, S. J. Bakker, R. P. Dullaart et al, 2014b Pleiotropic effects of lipid genes on plasma glucose, HbA1c, and HOMA-IR levels. Diabetes 63: 3149–3158.

Li, Q., Y. Li, Z. Zhang, T. R. Gilbert, A. H. Matsumoto et al, 2008 Quantitative trait locus analysis of carotid atherosclerosis in an intercross between C57BL/6 and C3H apolipoprotein E-deficient mice. Stroke 39: 166–173.

Liu, S., J. Li, M. H. Chen, Z. Liu and W. Shi, 2015 Variation in type 2 diabetes-related phenotypes among apolipoprotein E-deficient mouse strains. PLoS One 10: e0120935.

Lyons, M. A., R. Korstanje, R. Li, K. A. Walsh, G. A. Churchill et al, 2004 Genetic contributors to lipoprotein cholesterol levels in an intercross of 129S1/SvImJ and RIIIS/J inbred mice. Physiol. Genomics 17: 114–121.

Lyons, M. A., H. Wittenburg, R. Li, K. A. Walsh, G. A. Churchill et al, 2003 Quantitative trait loci that determine lipoprotein cholesterol levels in DBA/2J and CAST/Ei inbred mice. J. Lipid Res. 44: 953–967.

Lyons, M. A., H. Wittenburg, R. Li, K. A. Walsh, R. Korstanje et al, 2004 Quantitative trait loci that determine lipoprotein cholesterol levels in an intercross of 129S1/SvImJ and CAST/Ei inbred mice. Physiol. Genomics 17: 60–68.

Mani, A., J. Radhakrishnan, H. Wang, A. Mani, M. A. Mani et al, 2007 LRP6 mutation in a family with early coronary disease and metabolic risk factors. Science 315: 1278–1282.

Manning, A. K., M. F. Hivert, R. A. Scott, J. L. Grimsby, N. Bouatia-Naji et al, 2012 A genome-wide approach accounting for body mass index identifies genetic variants influencing fasting glycemic traits and insulin resistance. Nat. Genet. 44: 659–669.

Purcell-Huynh, D. A., A. Weinreb, L. W. Castellani, M. Mehrabian, M. H. Doolittle et al, 1995 Genetic factors in lipoprotein metabolism. analysis of a genetic cross between inbred mouse strains NZB/BINJ and SM/J using a complete linkage map approach. J. Clin. Invest. 96: 1845– 1858.

Rowlan, J. S., Z. Zhang, Q. Wang, Y. Fang and W. Shi, 2013a New quantitative trait loci for carotid atherosclerosis identified in an intercross derived from apolipoprotein E-deficient mouse strains. Physiol. Genomics.

Rowlan, J. S., Q. Li, A. Manichaikul, Q. Wang, A. H. Matsumoto et al, 2013b Atherosclerosis susceptibility loci identified in an extremely atherosclerosis-resistant mouse strain. J. Am. Heart Assoc. 2: e000260.

Rutti, S., J. A. Ehses, R. A. Sibler, R. Prazak, L. Rohrer et al, 2009 Low- and high-density lipoproteins modulate function, apoptosis, and proliferation of primary human and murine pancreatic beta-cells. Endocrinology 150: 4521–4530.

Saleheen, D., A. Nazir, S. Khanum, S. R. Haider and P. M. Frossard, 2006 R1615P: A novel mutation in ABCA1 associated with low levels of HDL and type II diabetes mellitus. Int. J. Cardiol. 110: 259–260.

Sehayek, E., E. M. Duncan, H. J. Yu, L. Petukhova and J. L. Breslow, 2003 Loci controlling plasma non-HDL and HDL cholesterol levels in a C57BL /6J x CASA /Rk intercross. J. Lipid Res. 44: 1744–1750.

Seidelmann, S. B., C. De Luca, R. L. Leibel, J. L. Breslow, A. R. Tall et al, 2005 Quantitative trait locus mapping of genetic modifiers of metabolic syndrome and atherosclerosis in low-density lipoprotein receptor-deficient mice: Identification of a locus for metabolic syndrome and increased atherosclerosis on chromosome 4. Arterioscler. Thromb. Vasc. Biol. 25: 204–210.

Shepherd, M., I. Ellis, A. M. Ahmad, P. J. Todd, D. Bowen-Jones et al, 2001 Predictive genetic testing in maturity-onset diabetes of the young (MODY). Diabet. Med. 18: 417–421.

Shi, W., N. J. Wang, D. M. Shih, V. Z. Sun, X. Wang et al, 2000 Determinants of atherosclerosis susceptibility in the C3H and C57BL/6 mouse model: Evidence for involvement of endothelial cells but not blood cells or cholesterol metabolism. Circ. Res. 86: 1078–1084.

Soranzo, N., S. Sanna, E. Wheeler, C. Gieger, D. Radke et al, 2010 Common variants at 10 genomic loci influence hemoglobin A(1)(C) levels via glycemic and nonglycemic pathways. Diabetes 59: 3229–3239.

Stylianou, I. M., S. R. Langley, K. Walsh, Y. Chen, C. Revenu et al, 2008 Differences in DBA/1J and DBA/2J reveal lipid QTL genes. J. Lipid Res. 49: 2402–2413.

Su, Z., X. Wang, S. W. Tsaih, A. Zhang, A. Cox et al, 2009 Genetic basis of HDL variation in 129/SvImJ and C57BL/6J mice: Importance of testing candidate genes in targeted mutant mice. J. Lipid Res. 50: 116–125.

Su, Z., Y. Li, J. C. James, A. H. Matsumoto, G. A. Helm et al, 2006a Genetic linkage of hyperglycemia, body weight and serum amyloid-P in an intercross between C57BL/6 and C3H apolipoprotein E-deficient mice. Hum. Mol. Genet. 15: 1650–1658.

Su, Z., Y. Li, J. C. James, M. McDuffie, A. H. Matsumoto et al, 2006b Quantitative trait locus analysis of atherosclerosis in an intercross between C57BL/6 and C3H mice carrying the mutant apolipoprotein E gene. Genetics 172: 1799–1807.

Teslovich, T. M., K. Musunuru, A. V. Smith, A. C. Edmondson, I. M. Stylianou et al, 2010 Biological, clinical and population relevance of 95 loci for blood lipids. Nature 466: 707–713.

Tian, J., H. Pei, J. C. James, Y. Li, A. H. Matsumoto et al, 2005 Circulating adhesion molecules in apoE-deficient mouse strains with different atherosclerosis susceptibility. Biochem. Biophys. Res. Commun. 329: 1102–1107.

Visscher, P. M., R. Thompson and C. S. Haley, 1996 Confidence intervals in QTL mapping by bootstrapping. Genetics 143: 1013–1020.

Wang, X., and B. Paigen, 2005 Genetics of variation in HDL cholesterol in humans and mice. Circ. Res. 96: 27–42.

Wang, X., R. Korstanje, D. Higgins and B. Paigen, 2004 Haplotype analysis in multiple crosses to identify a QTL gene. Genome Res. 14: 1767–1772.

Willer, C. J., and K. L. Mohlke, 2012 Finding genes and variants for lipid levels after genome-wide association analysis. Curr. Opin. Lipidol. 23: 98–103.

Wilson, P. W., W. B. Kannel and K. M. Anderson, 1985 Lipids, glucose intolerance and vascular disease: The framingham study. Monogr. Atheroscler. 13: 1–11.

Wilson, P. W., J. B. Meigs, L. Sullivan, C. S. Fox, D. M. Nathan et al, 2007 Prediction of incident diabetes mellitus in middle-aged adults: The framingham offspring study. Arch. Intern. Med. 167: 1068–1074.

Wittenburg, H., M. A. Lyons, R. Li, U. Kurtz, X. Wang et al, 2006 QTL mapping for genetic determinants of lipoprotein cholesterol levels in combined crosses of inbred mouse strains. J. Lipid Res. 47: 1780–1790.

Yaguchi, H., K. Togawa, M. Moritani and M. Itakura, 2005 Identification of candidate genes in the type 2 diabetes modifier locus using expression QTL. Genomics 85: 591–599.

Yuan, Z., H. Pei, D. J. Roberts, Z. Zhang, J. S. Rowlan et al, 2009 Quantitative trait locus analysis of neointimal formation in an intercross between C57BL/6 and C3H/HeJ apolipoprotein E-deficient mice. Circ. Cardiovasc. Genet. 2: 220–228.

Zhang, Y., B. Kundu, M. Zhong, T. Huang, J. Li et al, 2015 PET imaging detection of macrophages with a formyl peptide receptor antagonist. Nucl. Med. Biol. 42: 381–386.

Zhang, Z., J. S. Rowlan, Q. Wang and W. Shi, 2012 Genetic analysis of atherosclerosis and glucose homeostasis in an intercross between C57BL/6 and BALB/cJ apolipoprotein E-deficient mice. Circ. Cardiovasc. Genet. 5: 190–201.

Zhou, W., M. H. Chen and W. Shi, 2015 Influence of phthalates on glucose homeostasis and atherosclerosis in hyperlipidemic mice. BMC Endocr Disord. 15: 13-015–0015-4.

